# EEG error-related potentials encode magnitude of errors and individual perceptual thresholds

**DOI:** 10.1101/2022.12.06.519418

**Authors:** Fumiaki Iwane, Iñaki Iturrate, Ricardo Chavarriaga, José del R. Millán

## Abstract

Error-related potentials (ErrP) are a prominent electroencephalogram (EEG) correlate of performance monitoring, and so crucial for learning and adapting our behavior. Although there exists an agreement that ErrP signal awareness to errors, it remains poorly understood whether they encode further information. Here we report an experiment with sixteen participants during three recording sessions in which occasional visuomotor rotations of varying magnitude occurred during a cursor reaching task. We designed a brain-computer interface (BCI) to detect ErrP in single trials that provided real-time feedback to participants by changing the color of the cursor upon ErrP detection. The individual ErrP-BCI decoders exhibited good transfer across recording sessions and scalability over the varying magnitude of errors. Our results indicate that ErrPs encode not only the conscious perception of errors, but also their magnitude, in their amplitude and latency. Furthermore, a non-linear relationship between the ErrP-BCI output and the magnitude of errors predicts individual perceptual thresholds to detect rotations. The uncovered relationship is consistent with non-human primate studies, which found a similar relationship between the size of errors and simple spike activity of Purkinje cells, and we conjecture a cerebellar contribution to ErrP. Our experimental setup and findings open new avenues to probe and extend current theories of performance monitoring, which are based on response conflict tasks, by incorporating continuous human-interaction tasks as well as analysis of the ErrP complex as a whole rather than individual peaks.

## Introduction

Skillful tennis players can hit and return a ball with the desired effect because of their ability to identify errors between the predicted next states and the actual ones along the trajectory, which enables them to adjust their actions seamlessly (1). However, this is a challenging task for novice players. This individual variability in our ability to detect errors of varying degrees is key for sensorimotor and adaptation learning (2, 3). While previous research has uncovered a neural correlate of performance monitoring in the human electroencephalogram (EEG) elicited by the awareness of discrete erroneous actions, the so-called error-related potentials (ErrP) (4–8), little is known about whether these potentials encode the magnitude of errors. Some previous studies have reported co-varying characteristics of the ErrP with error magnitudes (9, 10) —such as phase coherence or amplitude—, while others did not (11). A major limitation of these studies is the use of a limited number of error categories (i.e., small, medium and large) that failed to capture a continuous relationship between the magnitude of errors and the characteristics of the ErrP. Such a relationship is critical to predict our individual ability to discern errors.

Here we designed an experiment to further investigate these two open questions –namely, does ErrP encode the magnitude of errors? Can they predict the individual threshold of error awareness?–, where occasional visuomotor rotations occur during a cursor reaching task. We also developed a brain-computer interface (BCI) to detect in real time the presence or absence of ErrP in the subjects’ EEG indicating a rotation of the joystick-to-cursor mapping. Critically, in our study induced errors are not constant, so as to avoid any adaptation process, and are variable in magnitude.

ErrPs are elicited when subjects perceive an erroneous action, either committed by themselves (4, 5) or by another person of agent (7, 12). The ErrP is characterized by two main deflections thought to be originated mainly from the anterior cingulate cortex (ACC) (6, 13–16), an initial error-related negativity (ERN) followed by a positive peak (Pe), observed over the frontocentral cortex. Further, a recent study found an increase of high-gamma band power along with ErrPs (17). Nevertheless, reports on the co-activating high-gamma activity are still limited and required further examination.

ErrP-BCIs have shown to enable seamless and intuitive interaction with external devices (8, 12, 18–23). Nevertheless, in these studies actions were executed at discrete steps, not continuously as required here. Some BCIs have succeeded in decoding the presence of an ErrP during a continuous task (24–27). However, these studies used experimental protocols where erroneous actions are abrupt stops of the device (24, 25), errors happen at the same moment during the interaction (26, 28), or the device fails to reach a target (27)—all extreme conditions that do not capture erroneous actions during a continuous interaction, but see Batzianoulis et al. (29). Furthermore, none of these previous studies characterized the ErrP signals as a function of the severity of the erroneous action.

We implemented a BCI system during the three recording sessions, in which ErrPs were elicited by a visuomotor rotation while the subject was using a joystick to continuously control a computer cursor to reach a target (Fig. 1a and b). Errors (rotations of the joystick-to-cursor mapping) were induced in 30% of the trials. On the first and second recording sessions, subjects performed the cursor reaching task with three different degrees of rotation; i.e., 20°, 40° and 60°. On the third session, subjects experienced a finer range of rotations; i.e., from 3° to 60° with a step of 3°, to characterize the ErrP over the continuous magnitude of errors. Furthermore, online continuous decoding of ErrPs were performed on the second and the third sessions in a “Plug-and-Play” manner (30) by using the data from the previous recording session. The BCI provided real-time feedback to participants by changing the color of the cursor upon ErrP detection.

**Fig. 1.**
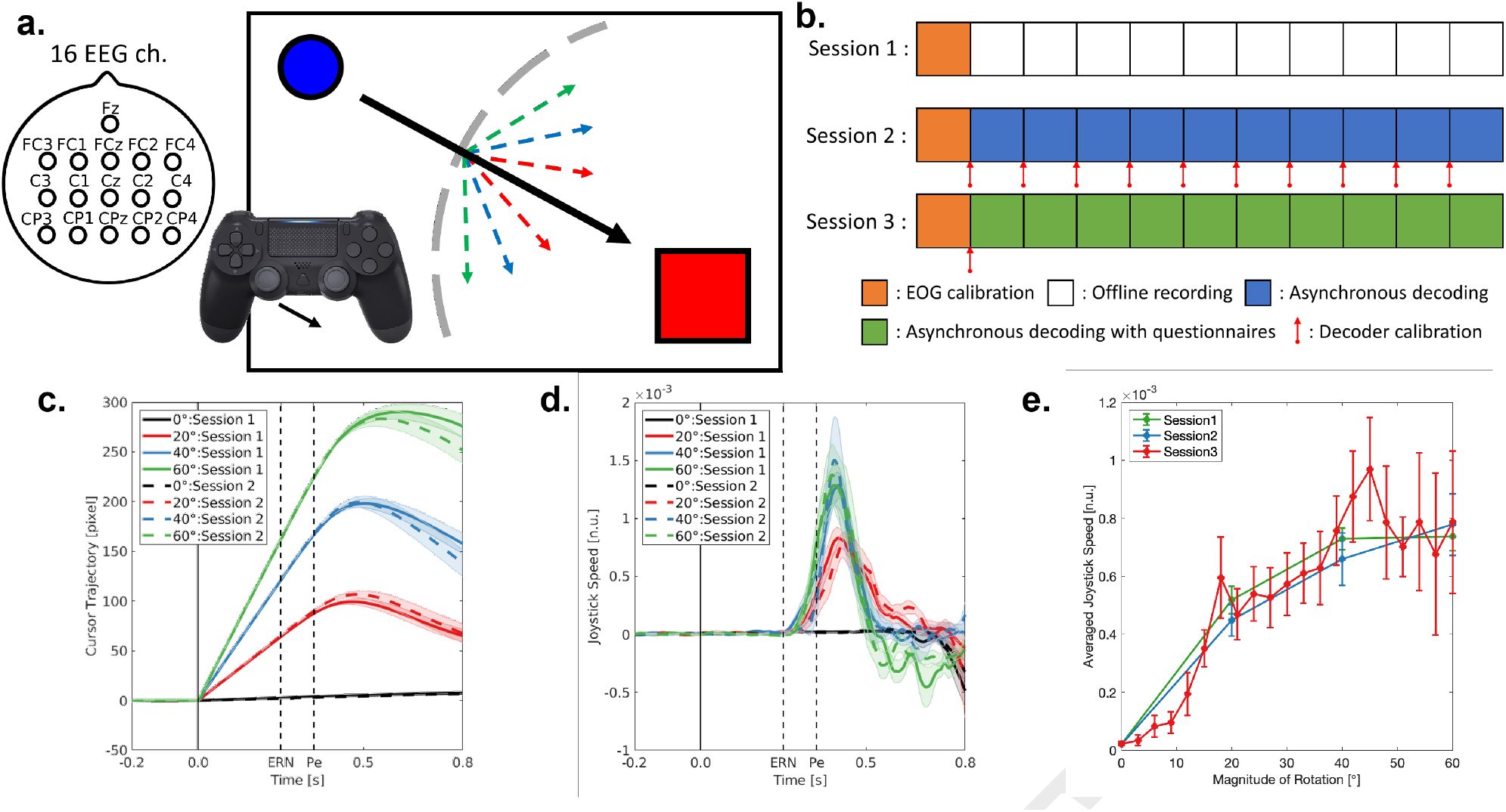
Experimental protocol and behavioral results. **a**. Overview of experimental protocol. The blue circle indicates a cursor controlled by the left-joystick of a game-pad. The red square represents the goal location where participants were instructed to bring the cursor as quickly as possible. Black arrow indicates the optimal cursor trajectory (straight line) to the goal. Gray dashed line represents the boundary to induce visuomotor rotations, which was randomly defined for each trial. In a correct trial (70% of trials), no visuomotor rotation occurred. In error trials (30%) of sessions 1 and 2, visuomotor rotations could be 20° (red), 40° (blue) or 60° (green), while in session 3 rotations were from 3° to 60° with a step of 3° (lines not shown). All lines were invisible to participants during the experiment. Note that participants cannot predict occurrence of visuomotor rotations not their magnitude. **b**. Experimental timeline for each recording session. 90 s of EOG calibration data was recorded before the cursor reaching task. In session 1, participants underwent the experiment without BCI feedback. In session 2, presence of ErrPs was continuously monitored while updating the decoder after each run. In session 3, continuous decoding was performed with a fixed decoder. **c., d**. Cursor deviations from straight trajectories and joystick speed in sessions 1 and 2 (mean, continuous or dotted lines, and standard error, shaded areas). **e**. Mean joystick speed of all sessions within the time window of [0.35, 0.45] s.

## Results

### Behavioral results

Participants generated similar cursor trajectories across recording sessions and consistently initiated corrective actions at 0.3 s with respect to the onset of the rotation (Fig. 1c and d). The peak joystick speed was observed at 0.41, 0.40 and 0.42 s for the first, second and third session, respectively. The mean joystick speed increased along magnitude of rotation while keeping consistency over the recording sessions (Fig. 1e). Variance of averaged joystick speed was larger on the third session compared with the first and the second session due to the lower number of trials recorded for each magnitude of rotation (n = 6 ± 1 (mean ± std)). Altogether, participants had a uniform behavior over the three recording sessions, and the speed of joystick movement was a function of the magnitude of errors.

### Time-frequency analysis

We observed band specific power modulation for theta [4, 8] Hz (26, 27), beta [18, 30] Hz (31, 32) and gamma [60, 90] Hz rhythms (17) (Fig. 2a and b). Theta band exhibited increased power modulation within the time window of [0.1, 0.6] s with respect to the onset of rotation. Beta and gamma band showed simultaneous modulations within the time window of [0.3, 0.8] s. While beta band power decreased, gamma band power increased when participants observed and corrected a visuomotor rotation (Wilcoxson’s signed-rank test, *p* < 0.001 for all three frequency bands). Theta and gamma band activity were modulated by the magnitude of rotation, while beta band activity remained stable for the different magnitudes of rotation (Fig. 2c). Furthermore, at ERN, we observed phase-amplitude coupling between high theta ([6, 8] Hz) and gamma band, while at Pe, coupling occurred between low theta ([4, 6] Hz) and gamma band (Fig. 2d). Similar to the temporal band power, phase-amplitude coupling increased for larger magnitudes of rotation. These results confirm that theta and gamma band activity, as well as their interaction, encode the magnitude of errors, while this is not the case for beta band activity.

**Fig. 2.**
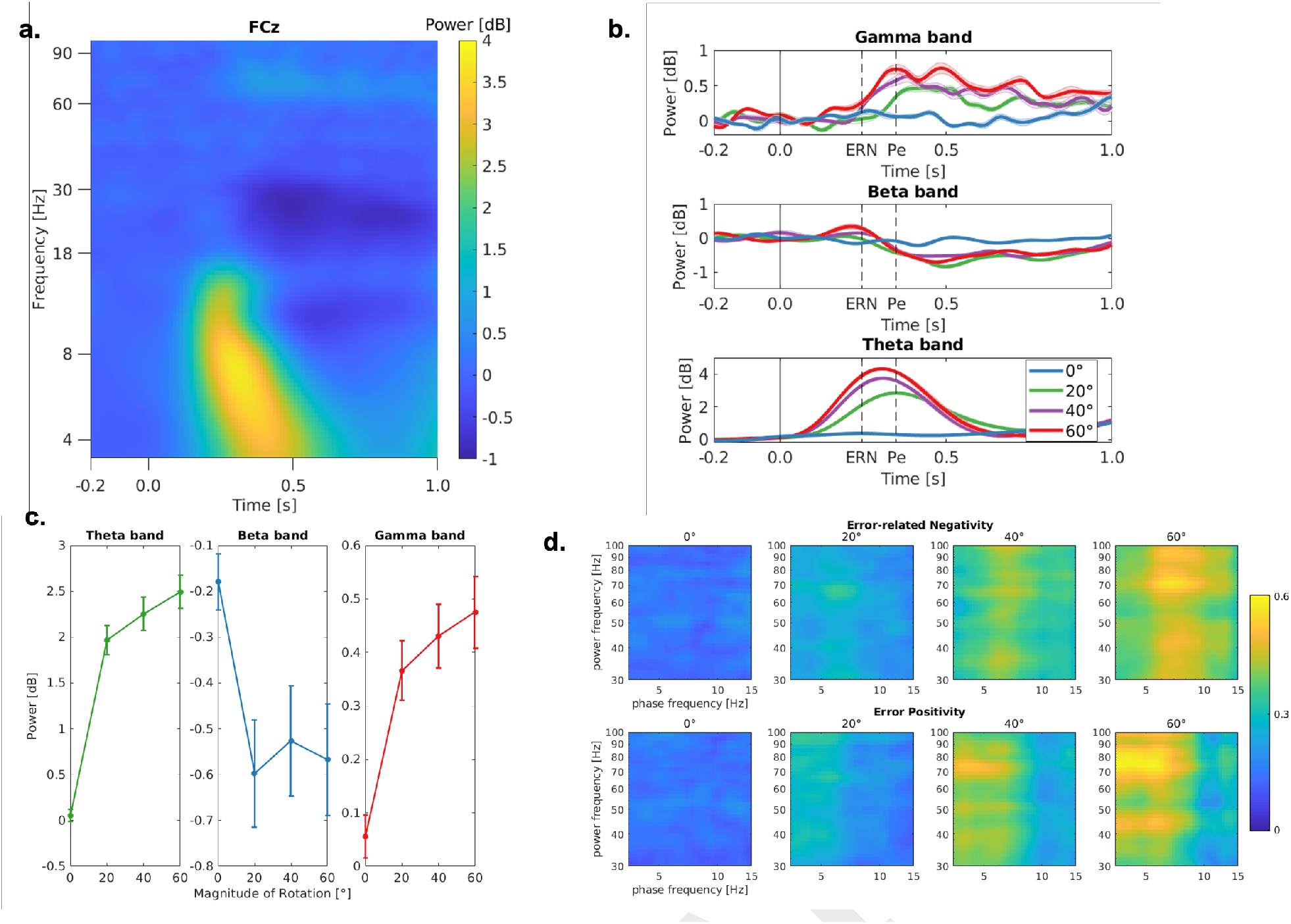
Time-frequency representations of ErrPs at FCz electrode on session 1 and 2. **a**. Time-frequency decomposition of ErrPs averaged over the all rotation trials with respect to the onset of rotation. Power was normalized by pre-stimuli baseline ([−0.25, 0.0] s). **b**. Theta (4-8 Hz), beta (18-30 Hz) and gamma band (60-90 Hz) power modulation with respect to the onset of event (*x* = 0 *s*) for each magnitude of rotation. Two dashed black lines at 0.25 s and 0.35 s indicate when ERN and Pe appeared. Theta band power increased within the time window of [0.1, 0.6] s with respect to the onset of rotation. Beta band power decreased, while gamma band power showed increase within the time window of [0.3, 0.8] s. **c**. Averaged band power modulation for each magnitude of rotation of session 1 and 2. Theta and gamma band power in erroneous trials were modulated as a function of magnitude of rotation (Pearson’s correlation analysis, Theta: *r* = 0.69, *p* < 0.001, Gamma: *r* = 0.39, *p* < 0.001), while beta band activity remained stable for the different magnitudes of rotation (Pearson’s correlation analysis, *r* = 0.09, *p* = 0.41). **d**. Theta-gamma phase-amplitude coupling at ERN and Pe. Note that levels of coupling is larger for trials with larger magnitude of rotations.

### BCI decoding results

The amplitude of ErrPs increased and the latency of the deflections shortened as the magnitude of rotations increased (Fig. 3a and b). In the third session (Fig. 3c) it is observed the two prominent deflections in the ErrPs when participants consciously perceived the rotation (*Yes*), and similar but delayed and smaller sequential peaks when participants were not sure whether they observed the rotation (*Maybe*). Topographical representations of all trials with rotation (insets in Fig. 3a, b and c) show similar patterns in the three recording sessions; namely, focal activation of the fronto-central area, especially at 0.35 s. These results suggest that ErrPs to encode magnitude of errors in its amplitude and latency and are stable across different recording sessions.

**Fig. 3.**
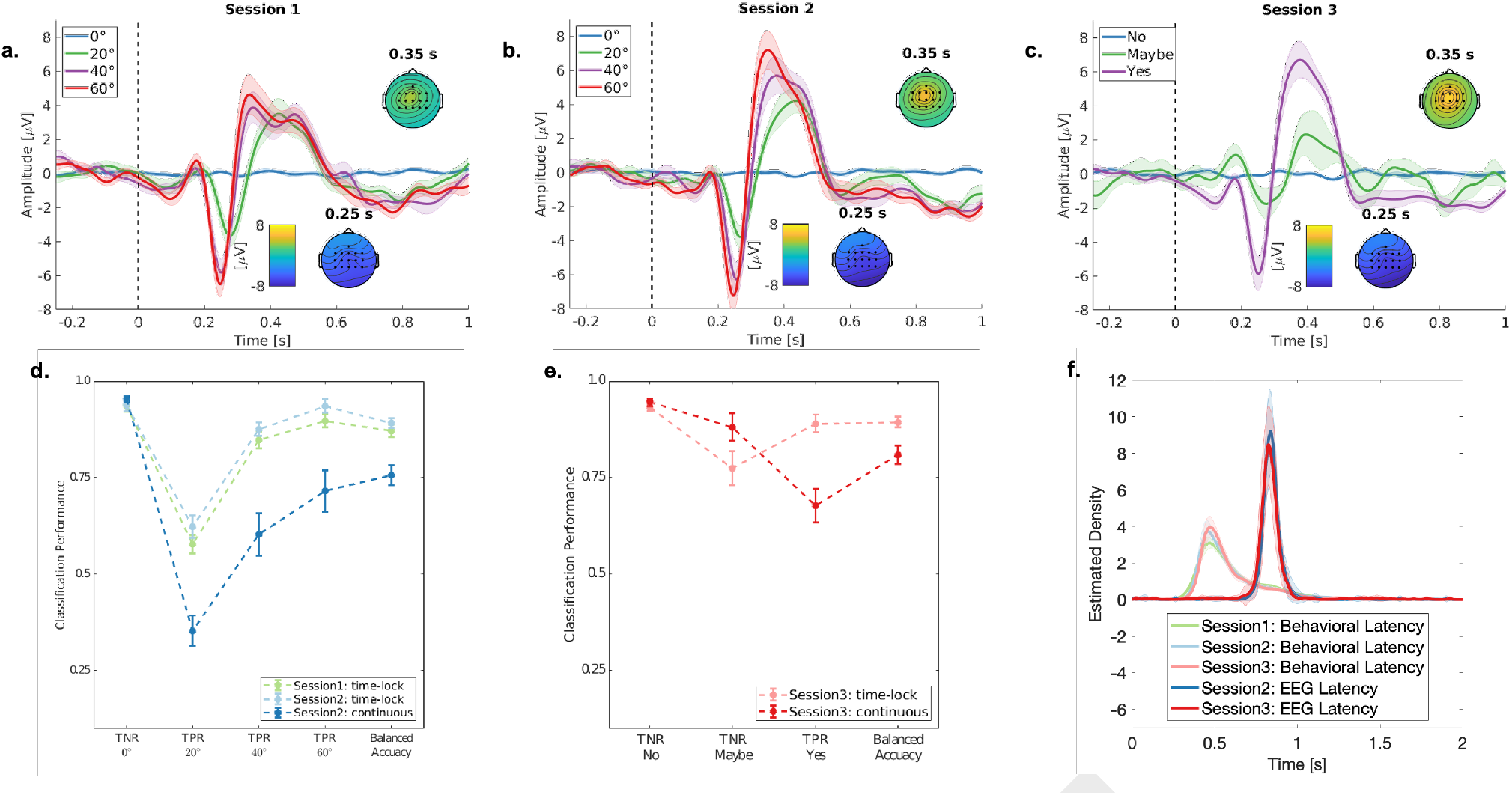
Grand-averaged signals and classification performance. **a., b., c**. Grand-averaged signals at FCz channel for each recording session, respectively. The dashed line (*x* = 0 *s*) represents the onset of rotation. On the first and the second recording session, each line corresponds to a magnitude of rotation; while lines correspond to the subjective behavioral answers on the third session. Each panel includes topographical representations averaged over trials with rotation of the negative and positive deflections at 0.25 and 0.35 s. **d., e**. Classification performance for time-locked classification and continuous decoding on the first two sessions and the third session, respectively. Note that True positive rate (TPR) increased as a function of magnitude on session 1 and 2.

Fig. 3d and e report both time-locked classification and online continuous decoding performance for each session (see also Table 1). Performance was similar across sessions (for different rotations in sessions 1 and 2, and balanced accuracy in the three sessions). TPR increased along with the magnitude of rotation in all conditions of classification (time-locked classification of the first and the second session and continuous classification of the second session) (Fig. 3d). The continuous classification performance (balanced accuracy) was significantly lower than the time-locked classification in the second and third sessions (Wilcoxon’s signed-rank test, *p* < 0.001 for both sessions), which illustrates the challenge to perform continuous decoding of ErrPs.

**Table 1.** Time-locked classification and online continuous decoding performance for each recording session (mean ± SE).

a Time-locked classification
| Training session | Test session | TNR of 0° | TPR of 20° | TPR of 40° | TPR of 60° | Balanced Accuracy |
| --- | --- | --- | --- | --- | --- | --- |
| Session 1 | Session 1 | 93.63 ± 1.54 | 57.73 ± 2.31 | 84.76 ± 2.13 | 89.74 ± 1.77 | 87.13 ± 1.65 |
| Session 1 | Session 2 | 93.84 ± 1.66 | 62.27 ± 2.88 | 87.55 ± 1.78 | 93.59 ± 1.77 | 89.16 ± 1.29 |
| Training session | Test session | TNR of No | TPR of Maybe | TPR of Yes |  | Balanced Accuracy |
| Session 1 and 2 | Session 3 | 93.13 ± 0.97 | 77.52 ± 4.38 | 89.99 ± 2.32 |  | 89.38 ± 1.38 |

b Continuous decoding
| Training session | Test session | TNR of 0° | TPR of 20° | TPR of 40° | TPR of 60° | Balanced Accuracy |
| --- | --- | --- | --- | --- | --- | --- |
| Session 1 | Session 2 | 95.30 ± 0.76 | 35.37 ± 3.86 | 60.37 ± 5.48 | 71.54 ± 5.34 | 75.64 ± 2.51 |
| Training session | Test session | TNR of No | TPR of Maybe | TPR of Yes |  | Balanced Accuracy |
| Session 1 and 2 | Session 3 | 94.54 ± 0.97 | 88.11 ± 3.52 | 67.84 ± 4.38 |  | 80.96 ± 2.36 |

During continuous decoding, presence of ErrPs were consistently detected with high temporal accuracy at around 0.850 s after the onset of rotation (Fig. 3f), which corresponds to the time window used to train the classifier, [0.2, 0.8] s (second session: 0.853 ± 0.01 s, third session: 0.846 ± 0.01 s). We then characterized the continuous relationship between the magnitude of rotations and the BCI output over the three sessions (Fig. 4). Estimated posterior probabilities of time-locked classification ranged from 0.21 (0 degrees) to 0.78 on the first two sessions and to 0.86 on the third session (60 degrees), while it ranged from 0.63 to 0.93 on the second and third sessions for continuous classification. For both, time-locked and continuous decoding, the estimated posterior probability linearly increased until 30°, then plateaued for larger rotations. Thus, an exponential model successfully captured the modulation over the magnitudes of visuomotor rotation (r-squared = 0.975 and 0.978, for time-lock and continuous classification, respectively). These results illustrate that the ErrP-BCI can not only detect errors, but also infer the magnitude of errors.

**Fig. 4.**
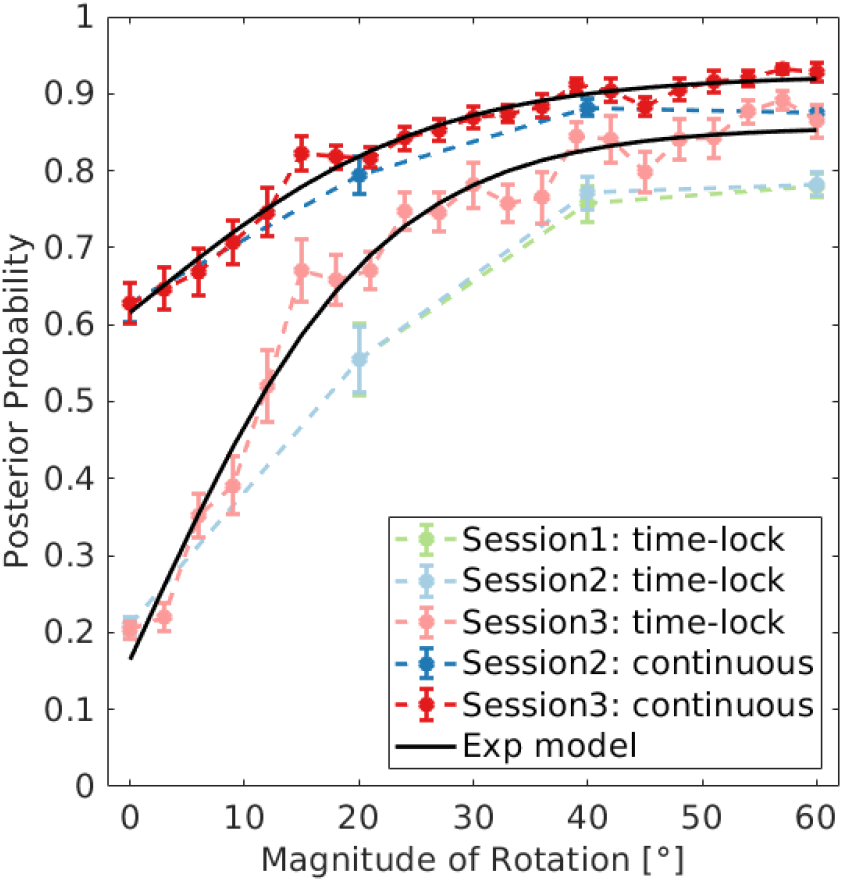
Estimated posterior probabilities of each recording session for each magnitude of rotation and fitted curve for both time-lock classification and continuous decoding on session 3. Note that estimated posterior probability is modelled as an exponential function of the magnitude of rotation.

### Inferring individual perceptual thresholds

Finally, we asked weather ErrPs predict individual perceptual thresholds to detect visuomotor rotations. To answer this question, we analyzed the ErrP-BCI output in the time-locked condition of the third session. The perceptual threshold is defined as the magnitude of rotation for which participants detected only 50% of occurrences. On average, participants started to answer “Maybe” at 8 ± 3 degrees and “Yes” at 12 ± 5 degrees of rotation (Fig. 5a). Most importantly, the behavioral answer and the output of the ErrP-BCI were modulated similarly, demonstrating the transferability of the decoder over sessions and the scalability of ErrPs over the different magnitudes of rotation. Fig. 5b represents the relationship between the behavioral individual perceptual threshold and the inferred individual perceptual threshold from the ErrP-BCI. Correlation analysis reveled a significant trend between the two percpetual thresholds (Spearman’s *r* = 0.61, *p* = 0.012), while the statistical test did not reveal significant difference between the two perceptual thresholds (Wilcoxon’s sign-rank test, *p* = 0.41). In summary, individual ability to detect errors was successfully inferred by characterizing the relationship between the magnitude of errors and their corresponding ErrPs.

**Fig. 5.**
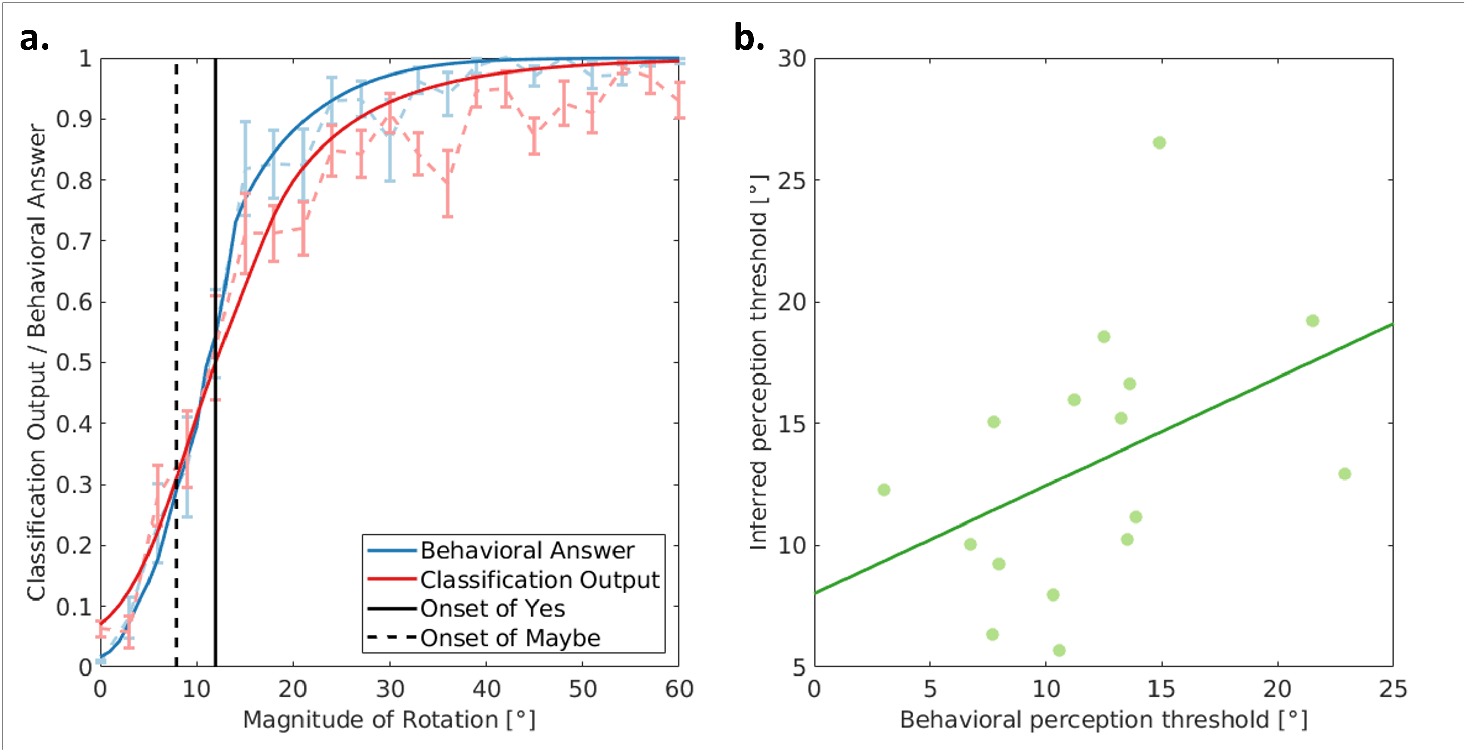
Inferring each subject’s individual perceptual threshold to detect a visuomotor rotation. **a**. Answer to the questionnaires collected after each trial and the results of the time-locked classification on the data collected on the third session. Black solid vertical line corresponds to onset of conscious perception (Yes) while black dashed line is onset of Maybe. For decoding output (red), 1 means that all trials were estimated as erroneous (there was a rotation), while 0 means that all trials were estimated as correct (no rotation). For behavioral answer (blue), 1 indicates that all trials of a given rotation was consciously perceived (i.e., Yes), while 0 means that all trials were not consciously perceived (i.e., Maybe or No). Behavioral answer and classification output increased similarly over the magnitude of rotation before and after the perceptual threshold (i.e., Onset of Yes). **b**. Scatter plot of the behavioral individual perceptual threshold and the inferred perceptual threshold based on the time-locked classification of ErrPs. Each dot corresponds to one subject. Green line corresponds to the regressed line by least-mean square.

## Discussion

Our results indicate that ErrPs encode not only the conscious perception of errors (i.e., visuomotor rotations in our experimental setup) (Fig. 3c), but also their magnitude, in their amplitude and latency (Fig. 3a and b). Furthermore, classification results from session 3 demonstrates how the ErrP decoder—built from data of sessions 1 and 2 that contains only rotations of 0°, 20°, 40° or 60°—generalizes across rotations ranging from 0° to 60° with a step of 3° and across recording sessions (Fig. 4). More fundamentally, analysis of the output of the ErrP-BCI shows that ErrPs predict individual perceptual thresholds to detect a visuomotor rotation.

As illustrated in Fig. 1a, participants used a joystick to control a cursor to reach a target. Unlike typical visuomotor rotation tasks (33–36), the visuomotor coupling between the cursor movement and the joystick was disrupted during the continuous cursor controlling task occasionally and with varying magnitudes in order to ensure participants could predict cursor trajectories and, so, elicit ErrPs in their EEG. This design enabled to induce stable and precise perturbations on the cursor trajectory and associated corrective actions across sessions (Fig. 1c and d). Furthermore, the elicited ErrPs were also stable across the three recording sessions (Fig. 3), in line with previous findings (8, 18, 37). On average, 46 and 30 days elapsed between consecutive sessions, respectively (Table 2).

**Table 2.** Time differences in days between the three recording sessions.

| Subject ID | s1 | s2 | s3 | s4 | s5 | s6 | s7 | s8 | s9 | s10 | s11 | s12 | s13 | s14 | s15 | s16 | Mean ± SE |
| --- | --- | --- | --- | --- | --- | --- | --- | --- | --- | --- | --- | --- | --- | --- | --- | --- | --- |
| Session1 - Session 2 | 52 | 63 | 42 | 42 | 51 | 49 | 56 | 56 | 51 | 59 | 43 | 44 | 35 | 24 | 35 | 37 | 46 ± 3 |
| Session2 - Session 3 | 25 | 13 | 40 | 35 | 38 | 38 | 28 | 31 | 26 | 11 | 32 | 45 | 52 | 32 | 21 | 10 | 30 ± 3 |

While most theories of performance monitoring consider ErrPs as a detector (38, 39), some studies found that ErrPs covary with the magnitude of errors (9, 10). However, these studies were limited to a short number of discrete error categories (i.e., small, medium and large), thus not capturing a continuous relationship between the two —which is crucial to explain our individual ability to detect errors. Moreover, whether or not ErrPs encode magnitude of errors has remained controversial, as other studies reported that different magnitudes of a visuomotor rotation do not affect ErrPs by comparing rotations of 45° and 180° (11). Our results extend and reconcile previous research as we have identified a continuous non-linear relationship between the magnitude of errors and the output of the ErrP-BCI, which starts plateauing after 30° of rotation (Fig. 4).

Having established this non-linear relationship, one fundamental question arises; namely, “Can we infer participants’ perceptual threshold to detect errors?”. As shown in Fig. 5a, behavioral answers to conscious perception of errors and inferred subjects’ responses from ErrPs were similarly modulated along with the magnitude of rotation. Interestingly, we observed that even at the magnitude where participants were unsure about the presence of the rotation (i.e., “Maybe”) the behavioral and inferred answers were closely aligned to each other. ErrPs for “Maybe” exhibited smaller and delayed deflections than for ErrPs elicited by conscious perception of the rotation (i.e., “Yes”, Fig. 3c). Additionally, Fig. 5b revealed a significant correlation between the behavioral perceptual threshold, or inflection point, and the inferred perceptual threshold. Taken together, our results suggests that ErrPs encode not only the magnitude of errors, but also the individual perceptual threshold to discern them consciously.

Our individual ability to detect errors of varying magnitudes is key to learn in motor adaptation tasks, where the cerebellum is known to play a major role (40–42). We speculate that neural correlates of error severity originate from Purkinje cells in the cerebellum and are projected onto the ACC via a cingulocerebellar circuit (43–45) to elicit ErrPs. In fact, the non-linear relationship between the magnitude of errors and the ErrP is consistent with non-human primate studies (46), in which monkeys manually tracked a pseudo-randomly moving target. Authors also found a non-linear relationship between size of errors and simple spike activity of Purkinje cells. Furthermore, patient studies seem to support that the cerebellum is involved in error processing (as reviewed in (47)) and there exists some evidence of cerebellar encoding of sensory prediction errors during motor tasks (48). Our experimental setup and findings open new avenues to probe and extend current theories of performance monitoring, which are based on response conflict tasks such as flanker (see, e.g., (38, 39) for reviews), by incorporating continuous human-interaction tasks as well as analysis of the ErrP complex as a whole rather than individual peaks. Moreover, while these theories interpret that the frontocentral Pe is simply associated with fast orienting and attentional processes, our results show Pe encodes qualitatively and quantitatively more information. In particular, while we observe a larger subsequent centropaerietal Pe at 400 ms (Fig S1), which has been proposed to drive adaptation (39), the two positive deflections reflect similarly the magnitude of errors.

Our results are not limited to the theta band activity of EEG. Völker et al. (17) recently reported the presence of the high-gamma band modulation during an Erikson flanker task in both invasive and noninvasive EEG recordings. High-gamma band ([60, 90] Hz) power increased when humans performed erroneous motor actions time-locked to the Pe component of ErrPs. Consistent with their findings, our results also demonstrate that the peak high-gamma power aligns with the Pe deflection in the same high-gamma band ([60, 90] Hz, Fig. 2b). Extending their results, Fig. 2d illustrates an increase only in theta and gamma band power, but not in beta. Furthermore, we observed a theta-gamma phase-amplitude coupling, which has often been observed during sensory signal detection and visual perception tasks (49, 50) (Fig. 2c). Although we have further analyzed datasets collected during our previous studies (27, 29), we did not observe such gamma band activity nor theta-gamma phase-amplitude coupling. A distinction between those previous studies and the present study is that in the latter subjects performed corrective actions upon perception of an error. Indeed, Krigolson *et al*. (51) showed that “correctability” of the task influences theta band activity; however, how the “correctability” modulates gamma band activity remains unclear. Altogether, these results suggest that theta-gamma coupling may occur only in the case where corrective actions are necessary to accomplish the task and covaries with magnitude of the perceived errors.

This study also uncovered a tight temporal relationship between corrective actions and the two characteristic electrophysiological deflections of ErrPs; namely, ERN and Pe. As shown in Fig. 3, we observed ERN at 0.25 s with respect to the rotation onset, which corresponds to the time period where participants initiate the corrective actions to re-aim the target (Fig. 1d). Similarly, Pe was observed at 0.35 s, in the middle of corrective actions. Interestingly, temporal patterns of both joystick speed and high-gamma band power were monotonically increasing from baseline towards its peak value within the time window between ERN and Pe. Overall, our results suggest presence of rapid information transfer between theta and high-gamma band activity to activate corrective actions of right amount after detection of an error.

Finally, our study has implications for single-trial decoding of ErrPs during continuous human-computer interaction. The individual BCI decoders exhibited good transfer across recording sessions and scalability over the varying magnitude of errors. As shown in Fig. 3, stable ErrP patterns over the recording sessions enabled us to deploy “Plug-and-Play” ErrP-BCIs on the second and the third session, in which online decoding of ErrPs were carried out since the first run of the session by transferring the decoder from the previous recordings. Transfer learning to detect ErrPs has been investigated across different error types (52, 53), cognitive workload conditions (27, 54) and participants (25, 55); however, they often reported degraded classification performance. This was not the case in our experiments, where we observed consistent classification performance by transferring the personalized decoder over the different decoding sessions, especially during online continuous classification (Fig 3d and e, and Table 1). Nevertheless, our results revealed the difficulty to detect small errors compared to large ones due to the reduced amplitude and delayed deflections of their elicited ErrPs (Fig. 3a and b, and Fig. 4). Importantly, the latency to detect ErrPs was close to the optimal latency and very stable across sessions during online continuous decoding (Fig. 3f), thus demonstrating the temporal precision of the continuous ErrP-BCI. We conjecture that such a kind of ErrP-BCI could enhance operation of assistive devices by people with severe motor disabilities. Indeed, because of their degraded residual control, these subjects will occasionally deliver wrong commands similar to the visuomotor rotations in our experiments, and will not be fast enough to execute corrective actions before the output of the ErrP-BCI will be available.

## Materials and Methods

### Participants

Sixteen able-bodied healthy subjects (four female, 23 ± 1 years old) participated in the experiment, which consisted of three recording sessions (days). The experimental protocol was approved by the local ethics commission (PB_2017-00295). Written informed consent was collected from all participants before conducting the experiment. During the experiment, participants sat on a comfortable chair in front of a laptop with a 14-inch display that visualized the experimental protocol.

### Experimental design

Fig. 1a illustrates the experimental protocol. A blue circle represented the cursor which was controlled by the left joystick of a game-pad (DualShock4, Sony, Japan), and a red square represented the goal location where participants were instructed to bring the cursor as quickly as possible. The speed of the cursor was kept constant (750 pixel/s) during the cursor reaching task as long as the joystick was pressed. Averaged duration of a trial was 3.5 ± 0.2 s. The goal location was randomly chosen for each trial among six potential locations; i.e., top-left, middle-left, low-left, top-right, middle-right and low-right. Initial position of the cursor was at the other side of the target at a random height to ensure enough distance (at least 1960 pixel) between the initial position of the cursor and the goal. For example, if the goal was on the right side, the cursor was placed on the left side at a random height and vice versa. In 30% of trials, the joystick-to-cursor mapping was rotated when the cursor exceeded a pre-defined invisible boundary. The boundary was randomly determined for each trial. Trials with a visuomotor rotation were defined as “erroneous trials”, and others as “correct trials”. The magnitude of rotation was fixed to 20°, 40° and 60° on the first and the second sessions. On the third session, the magnitude of rotation was between 3° to 60° with a step of 3°, and after each trial participants were asked whether they perceived the rotation, maybe or not. Table 2 reports the days difference between each recording session for each subject. As illustrated in Fig. 1b subjects performed 10 runs of 40 trials in each session (400 × 3 trials in total). Before these runs, participants made eye movements for 90 s for estimating regression parameters to remove EOG artifacts from their EEG during the actual experiment (56).

Participants executed the experiment without receiving online feedback on the first session. We provided subjects with online feedback on the second and the third sessions by changing the color of the cursor upon ErrP detection. On the second session, presence of ErrPs was continuously monitored during cursor control, and the decoder was re-calibrated after each run. On the third session, we carried out continuous decoding of ErrPs without re-calibrating the decoder, which was built with data from the first and the second sessions, except for the parameters to remove EOG artifact.

### EEG and EOG acquisition

16 EEG and 3 EOG electrodes were recorded throughout the experiment (two synchronized g.USBAmp, g.tec medical engineering, Austria). EEG electrodes were located at Fz, FC3, FC1, FCz, FC2, FC4, C3, C1, Cz, C2, C4, CP3, CP1, CPz, CP2 and CP4 in 10/10 international coordinates, and EOG electrodes were placed at above the nasion and below the outer canthi of the eyes. The ground electrode was placed on the forehead (AFz) and the reference electrode was placed on the left earlobe. We used the same reference and ground electrode for both EEG and EOG signals. EEG and EOG signals were notch filtered at 50 Hz by the amplifiers. To reduce signal contamination, participants were asked to avoid excessive eye movements and blinks during trials.

### Time-frequency analysis

To characterize the time-frequency representation of ErrPs elicited by visuomotor rotations, we computed a discrete wavelet time-frequency decomposition (57). After applying a 2nd order non-causal high-pass Buttterworth filter with the cut-off frequency of 1 Hz and current source density (CSD), EEG signals were epoched in the time window [−0.2, 1.0] s with respect to the onset of rotations. For each single-trial EEG epoch we performed the Morlet Wavelet time-frequency decomposition (58) in the frequency range [3, 100] Hz, resulting in a wavelet coefficient matrix with 126 time points and 97 log-spaced frequency bins. Each of time windows were composed of 427 samples (0.83 s) overlapped by 51 samples (0.1 s); while the number of cycles ranges from 3 to 20 at highest. The resulting coefficients were used to extract spectral power and phase-amplitude coupling.

To compute the spectral power induced by a visuomotor rotation we used the event-related spectral perturbation (ERSP) approach (59). This approach is less sensitive to noisy trials than classical baseline correction methods, and produces a non-skewed power distribution. In detail, separately for each subject and experimental condition, we apply a single-trial full-epoch baseline correction, before averaging across trials and removing the trial-averaged pre-stimulus (i.e., from −0.25 to 0.0 s) baseline. Baselines were corrected using the gain model assumption (i.e., divide by the baseline) instead of the additive model (i.e., subtract the baseline) (59, 60). Finally, the trial-averaged pre-stimulus corrected time frequency coefficients were log-transformed (10log_10_).

To compute the phase-amplitude coupling, we first identified the latency of ERN and Pe (0.25 and 0.35 ms, respectively), then used the mean vector length (MVL) method. MVL has been reported to be more suitable for high signal-to-noise ratio data as it is more sensitive to coupling strength and width compared to other methods such as modulation index or phase-locking value (61). This method estimates the coupling between phase frequency *f_p_* and amplitude frequency *f_a_* from a number of epochs *N*, by mapping phase time series 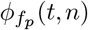 and amplitude time series 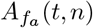 to a complex-values vector at each time point, *t*, and each epoch, *n*. To quantify the coupling between *f_p_* and *f_a_*, the MVL method measures the length of the average vector and computes phase amplitude coupling as follows:

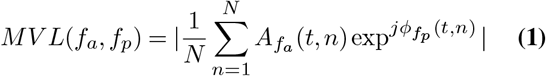

### BCI Decoding analysis

EEG signals were band-pass filtered with a 4th order causal Butterworth filter with cutoff frequencies of [1, 10] Hz, for both online and offline analyses.

To build the individual decoders, ErrPs were segmented into epochs in the time window of [0.2, 0.8] s with respect to the onset of the visuomotor rotation. Firstly, to enhance the signal-to-noise ratio of the EEG, we applied a spatial filter based on canonical correlation analysis (CCA) (37, 62). This spatial filter method transforms the averaged ErrPs to a sub-space containing different ERP components (63). Only the first three components were kept for the subsequent analysis. This number was determined based on the data collected on the first session by performing pseudo-continuous decoding in cross validation. For every trial, we extracted three complementary types of features: the decimated signal amplitude per CCA component at 64 Hz; power spectral densities per CCA component from 4 to 10 Hz with a step of 2 Hz; and the covariance matrix on Riemannian geometry, which computes a low-dimensionality manifold representation from a non-linear combination of the EEG component space (64). In order to include information of the EEG temporal dynamics in the Riemannian spatial covariance matrix, the epoch *X* was augmented with an individual template *T* representing the grand average of erroneous trials in the training set:

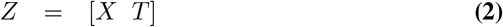

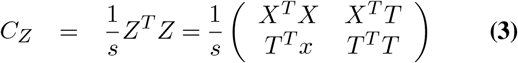

where *s* denotes number of time samples of an epoch. The covariance between *X* and *T* allows to capture the temporal dynamics of multi-component EEG signals with respect to the template. The covariance matrix was then projected on the tangent space, computed only on the training dataset (65). All computed features were concatenated and normalized within the range of [0, 1]. From this feature vector **x**, we computed the estimated posterior probability of having detected an error, *p*(*error*|**x**) using diagonal linear discriminant analysis (LDA):

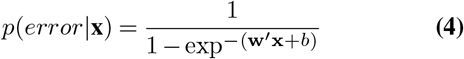

In order to continuously monitor the presence or absence of ErrPs, we analyzed the EEG signal with a sliding window, online or offline (24, 27).

To compute the estimated posterior probability of the training data without overfitting, aforementioned signal processing was performed in a leave-one-run-out cross-validation manner. Training folds were used to create an ErrP decoder, while the testing fold was used to compute the continuous modulation of posterior probability during cursor control with a sliding window at 32 Hz, 16 sample shift. In order to infer participants’ erroneous perception in a continuous manner and provide them with the feedback upon the detection of ErrPs, two hyperparameters were optimized besides the ErrP decoder; namely, smoothing factor and decision threshold. The smoothing factor indicates length of the time window for an online moving average filter on the estimated posterior probabilities, ranging from 1 to 16 with a step of 1. The decision threshold determined presence or absence of erroneous perception, ranging from 0 to 1 with a step of 0.01. Hyperparameter optimization was carried out in a grid search manner. Specifically, we computed the Matthew’s correlation coefficient (MCC) for each pair of hyperparameters based on a confusion matrix. With a pair of hyperparameters, a correct trial was properly classified if the smoothed posterior probability did not exceed the decision threshold (True Negative). If the smoothed posterior probability in a correct trial exceeded the decision threshold, the trial was considered as erroneous (False Positive). On the other hand, an erroneous trial was correctly classified if the posterior probability exceeded the decision threshold in a time window of [0.5, 1.1] s with respect to the onset of the rotation (True Positive). This time window of [0.5, 1.1] s was determined as the optimal latency was 0.8 s and we allowed 0.3 s of temporal difference with respect to the optimal latency. If the averaged posterior probability in an erroneous trial exceeded the decision threshold outside the aforementioned time window or did not exceed the decision threshold, it was considered as wrong classification (False Negative). For each pair of hyperparameters we computed their MCC, a 16 × 101 matrix for each testing fold. The pair of hyperparameters with the highest MCC, averaged over the testing folds, was chosen as the optimal. Once the pair of optimal hyperparameters was determined, we used all the available data to re-compute the ErrP decoder to be deployed subsequently for online continuous decoding. Decoder re-calibrations during the second session—from runs 2 to 10—used the all available data up to that moment (i.e., first session plus previous runs in second session), and reestimated the pair of optimal hyperparameters. On the second and the third sessions, we provided participants with the online feedback when the smoothed posterior probability exceeded the decision threshold for the first time in a trial.

### Individual perception threshold

In order to model behavioral perception and estimated posterior probabilities over the magnitudes of rotation (Fig. 5), we used the following exponential function:

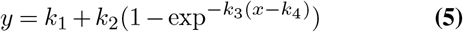

where *k*_1−4_ represent the initial value, maximum value, magnitude of slope and inflection point of the curve, respectively; and *x* represents the magnitude of rotation. Parameter *k*_1−4_ were estimated by gradient descent with the objective function defined as the root mean square error between the observation and the fitted model.

## ACKNOWLEDGEMENTS

This work was supported by the Hasler Foundation, Switzerland.

