## Supplemental Figure 1 for "EEG error-related potentials encode magnitude of errors and individual perceptual thresholds"

Supplementary Figures

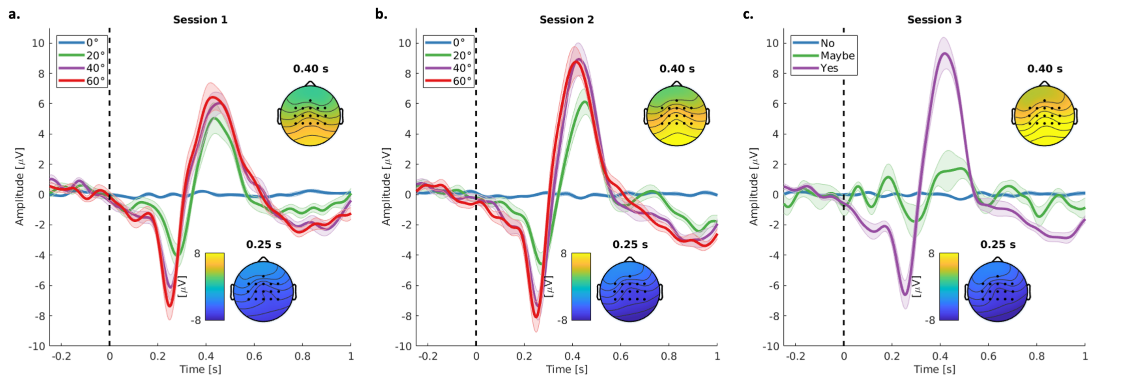


**Supplemetary Figure 1:** **a., b., c.** Grand-averaged signals at CPz channel for each recording session, respectively. The dashed line ($x = 0\ s$) represents the onset of rotation. On the first and the second recording session, each line corresponds to a magnitude of rotation; while lines correspond to the subjective behavioral answers on the third session. Each panel includes topographical representations averaged over trials with rotation of the negative and positive deflections at 0.25 and 0.40 s.
